# Comparative genomics reveals contraction in cytosolic glutathione transferase genes in cetaceans: implications for oxidative stress adaptation

**DOI:** 10.1101/485615

**Authors:** Ran Tian, Inge Seim, Wenhua Ren, Shixia Xu, Guang Yang

## Abstract

Cetaceans, a highly specialized group of aquatic mammals, experience oxidative stress induced by reactive oxygen species (ROS) production associated with the apnea/reoxygenation. The glutathione transferases (GST) family possesses multiple functions in detoxification and antioxidant defenses. However, the molecular evolution of GST family in cetaceans is still poorly investigated. Here, we characterized the GST gene family across 21 mammalian genomes including cetaceans and terrestrial relatives. Overall, 7 GST classes were identified, showing GSTs are ubiquitous and conservative to all mammals. Some of GSTs are lineage-specific duplication and loss, in line with a birth-and-death evolutionary model. We detected positive selection sites that possibly influence GST structure and function, suggesting adaptive evolution of GSTs is important for defending mammals from various types of noxious environmental compounds. There is evidence for loss of alpha and mu GST class in cetacean lineages when compared to their terrestrial relatives, consisting with the lower GST activities observed in cetaceans. Notably, we find that retained *GSTA1*, *GSTA4* and *GSTM1* in cetaceans, indicating a vital role in against lipid peroxidation and superoxide. Besides, variation in gene number, enzyme activity and selection pressure of GSTs between marine mammals suggests there is a divergent evolution of GSTs in aquatic species that might be associated to diving ability and oxidative status with different habitats. Summarily, our findings demonstrate that the GST family in cetaceans has been subject to evolutionary dynamics as response for their adaptations to oxidative stress, and highlighted the differential selection associated with different life history traits among mammals.

## Introduction

One of the classical examples of evolution is the return of terrestrial vertebrates to an aquatic environment – manifested by functional (secondary) adaptations in species whose ancestors departed an aquatic environment hundreds of millions of years earlier. The order cetacea (whales, dolphins, and porpoises) is a model group of air-breathing marine mammals that transitioned to an aquatic lifestyle at least three times independently 53 to 55 million years ago (Uhen 2007). The majority of cetaceans make shallow, short dives. This includes the common dolphin (*Delphinus delphis*), which usually dives down to 200 m and remains submerged for 5 minutes (Schreer and Kovacs 1997), however, a few cetaceans are capable of deep, long dives. For example, the sperm whale (*Physeter catodon*) can dive to a depth of 3,000 m and stay underwater for at least 138 minutes (Schreer and Kovacs 1997). Regardless of different diving abilities, all cetaceans face the tremendous challenge posed by a lack of oxygen associated with dives, so-called asphyxia (the integration of hypoxia, hypercapnia, and acidosis) (Elsner and Gooden 1983). In response, cetaceans have evolved morphological, physiological, and biochemical adaptations. This includes improved oxygen stores, achieved by elevated hemoglobin and myoglobin levels, and modifications in energy metabolism, including increased activity of glycolytic enzymes to strengthen tolerance to hypoxia (Ramirez et al. 2007) and amino acid changes in the rate-limiting enzymes of the gluconeogenesis pathway (Tian et al. 2017). A number of complex cardiovascular responses play a vital role in the conservation of oxygen in cetaceans; especially bradycardia and peripheral vasoconstriction which preferentially augment or maintain blood flow to the central nervous system and heart, but reduce flow in peripheral tissues such as kidney, liver, and skeletal muscle (Panneton 2013). Diving mammals (e.g. harbor seal) virtually ceases blood flow to the kidneys during diving, leading to a renal ischemia (Blix 2018). However, the kidneys of diving mammals have improved reperfusion capacity and urine production (Blix 2018). Thus, cetaceans can continuously alternate from hypoxic to normoxic environments, manifested as regional ischemia (blood flow restriction) during diving, and followed by prompt reperfusion at the water surface (Panneton 2013).

While diving is facilitated by improved hypoxia resistance and tolerance to repeated ischemia/reperfusion events, reoxygenation after a dive would be expected to result in the production of reactive oxygen species (ROS) (Li and Robert 2002).

These molecules are very reactive and unstable byproducts of oxygen metabolism and include superoxide anion (O_2_^-^) and hydrogen peroxide (H_2_O_2_) (Apel and Hirt 2004). ROS can oxidize all cellular macromolecules (DNA, proteins, and lipids), leading to oxidative damage (Apel and Hirt 2004). To prevent such damage cells employ antioxidant defense systems (AD) (Birben et al. 2012). Several studies on diving mammals have demonstrated that antioxidant enzymes are critical components of these systems. These include catalase (CAT), superoxide dismutase (SOD), glutathione-transferase (GST), glutathione reductase (GSR), and glutathione peroxidase (GPX) (Birben et al. 2012). For example, higher enzyme levels (glutathione disulfide, glutathione disulfide reductase, and glucose-6-phosphate dehydrogenase) in the glutathione (GSH) antioxidant defense system has been reported to be associated with protection against dive-induced ischemia/reperfusion in ringed seal (*Phoca hispida*) tissues (heart, kidney, lung, and muscle) (Vázquez-Medina et al. 2007). Tissues of diving mammals also have high levels of antioxidant enzymes like SOD, GSR, and GPX compared to terrestrial mammals (Vázquez-Medina et al. 2006; Wilhelm et al. 2002; Elsner et al. 1998; García-Castañeda et al. 2017). These data suggest that diving mammals have indeed evolved physiological adaptions in their antioxidant defense systems

One important group of antioxidant enzymes are GST, a large supergene family (Hayes et al. 2005). As enzymes in Phase II detoxification, acting downstream of the Phase I cytochrome P450 supergene family of enzymes expressed primarily by the liver, GST catalyses the conjugation of the GSH with reactive electrophilic compounds to form thioethers, called S-conjugates (Hayes et al. 2005). GSTs have evolved together with GSH and are abundant and widely distributed in animals, plants, insects, and microbes (Board and Menon 2013). They are also believed to protect tissues from endogenous organic hydroperoxides caused by oxidative stress (Board and Menon 2013). The mammalian GSTs have been divided into three superfamily classes which are structurally distinct and have separate evolutionary origins (Board and Menon 2013): cytosolic, mitochondrial, and microsomal transferases. Cytosolic GSTs represent the largest class and consists of seven distinct subclasses: alpha (encoded by human chr 6), mu (chr 1), theta (chr 22), pi (chr 11), zeta (chr 14), sigma (chr 4), and omega (chr 10). The number of genes in each subclass varies across the phylogenetic tree. For example, human alpha (*GSTA1* to *GSTA5*) and mu (*GSTM1* to *GSTM5*) have five enzymes each, omega (*GSTO1* and *GSTO2*) two members each, and zeta (*GSTZ*) has only one member (table S1). Conventionally, members within each GST subclass shares greater than 40% amino acid sequence identity, while members of different subclasses share less than 25% amino acid identity (Wu and Dong 2012). Although diverse mammals have retained each cytosolic GSTs subclass, there is variation in the number of genes in each subclass. For example, there is one pi subclass (GSTP) gene in humans and two pi subclass genes in mice (Bammler et al. 1994). There are now numerous manuscripts on the molecular evolution of GST genes in plants and animals (Ding et al. 2017; Low et al. 2007; Monticolo et al. 2017). For example, it has been suggested that modifying the number and diversity of GST genes has shaped the ecological adaptation of birds (Khan 2014). (Liu et al. 2015) reported that polyploidy-derived GST duplicated gene pairs might be beneficial for the defense responses of soybean. In contract, we are not aware of studies comparing GST genes in terrestrial and marine mammals. Here, we performed a comparative genomics analysis of cytosolic GST genes in 21 mammals, including 10 from the three distinct orders of aquatic mammals. We reveal putative genetic mechanisms for the adaptation of oxidative stress in cetaceans and improve the understanding of the genetic and evolutionary dynamics of GST superfamily in mammals.

## Methods and materials

### Sequence retrieval

We obtained all known human GST genes by perusing research articles and recent reviews (Board and Menon 2013; Morel et al. 2002) and by downloading coding sequences (CDS) from GenBank (http://www.ncbi.nlm.nih.gov). The corresponding GenBank accession numbers are listed in table S1. Employing human protein sequences as queries, BLASTn and tBLASTn searches were performed using an in-house Python script on local databases constructed from downloaded genomic sequences from 20 species. These included seven cetaceans: bottlenose dolphin (*Tursiops truncatus*), killer whale (*Orcinus orca*), Yangtze river dolphin (*Lipotes vexillifer*), Yangtze finless porpoise (*Neophocaena asiaeorientalis*), minke whale (*Balaena acutorostrata*), bowhead whale (*Balaena mysticetus*), sperm whale (*Physeter macrocephalus*); two pinnipeds: Weddell seal (*Leptonychotes weddellii*), Pacific walrus (*Odobenus rosmarus divergens*); one sirenian: Florida manatee (*Trichechus manatus latirostris*); and ten terrestrial mammals: cow (*Bos taurus*),

Tibetan yak (*Bos mutus*), sheep (*Ovis aries*), Tibetan antelope (*Pantholops hodgsonii*), dog (*Canis lupus familiaris*), horse (*Equus caballus*), microbat (*Myotis lucifugus*), mouse (*Mus musculus*), naked mole rat (*Heterocephalus glaber*), and elephant (*Loxodonta africana*). Selected genome sequencing and assembly information of each species are listed in table S2. We retrieved multiple hits for each query and identified non-redundant hits by extending 1,000 bp at both 5’ and 3’ ends. All hits were used to identify and annotate GST genes following the canonical gt/ag exon/intron junction rule (Cheng et al. 1995). In addition, the online resource GENEWISE (http://www.ebi.ac.uk/Tools/psa/genewise) (Birney et al. 2004) was employed to identify open reading frame (ORF) sequences. To enhance the comprehensiveness of our approach, we queried a species genome with its respective predicted GST sequences using BLAST. Mammalian GSTs were named according to the revised GST nomenclature of human (Nebert and Vasiliou 2004). All identified GST genes were categorized into three categories – based on amino acid composition, unique motifs, and BLAST and alignment results: 1) intact gene (putatively functional genes), a complete CDS region with the canonical structure typical of GST families; 2) partial gene, putative functional protein, but missing a start codon and/or stop codon; 3) pseudogene, highly similar to functional orthologs but with (a) inactivating mutation(s) and/or stop codon(s). Synteny analysis was also performed by retrieving the flanking genes for each GST classes using abundant BLAST searches and the Genomicus v93.01 (Nguyen et al. 2017).

### Phylogenetic analysis

The phylogenetic relationships of putative GST members in each subclass were estimated by maximum likelihood and Bayesian methods, as implemented in RAxML v8.0.26 (Stamatakis 2014) and MrBayes v3.1.2 (Ronquist and Huelsenbeck 2003). Nucleotide sequences were aligned using MUSCLE v3.6 (Edgar 2004), implemented in SeaView v4.5.4 (Gouy et al. 2009), and the manually corrected upon inspection. MrModeltest was used to estimate the best-fit model (SYM+G) of nucleotide substitution (Nylander 2009). For the RAxML analyses the ML phylogeny was estimated with 1,000 bootstrap replicates. For MrBayes analyses we performed two simultaneous independent runs for 50 million iterations of a Markov Chain, with six simultaneous chains, sampling every 1,000 generations. A consensus tree was obtained after discarding the first 25% trees as burn-in.

### Gene family analyses

To identify expanding and contracting gene ortholog groups across the mammalian phylogeny, we estimated the gene numbers on internal branches using a random birth and death process model implemented in the software CAFÉ v3.0, a tool for the statistical analysis of the evolution of the size of gene families (De Bie et al. 2006). We defined gene gain and loss by comparing cluster size differences between ancestors and each terminal branch among the phylogenetic tree. The ultrametric tree, based on the concatenated orthogroups, was estimated with BEAST v1.10 using Markov chain Monte Carlo (MCMC) with fossil calibrations and a Yule tree prior (Suchard et al. 2018). The fossil-derived timescales were searched from TimeTree (http://www.timetree.org). The analyses ran for 10 million generations, with a sample frequency of 1,000 and a burn-in of 10%.

### Adaptive evolution analyses

To evaluate the positive selection of all GST genes during mammalian evolution, codon substitution models implemented in the *codeml* program in PAML v4.4 were applied to GST gene alignments (Yang 2007). Two pairs of site-specific modes were tested comparatively using the likelihood ratio test (LRT): M8 (beta & ω) vs. M8a (beta & ω = 1) (Swanson et al. 2003). M8 estimates the beta-distribution for the ω and takes into account positively selected sites (ω > 1), with the neutral model M8a not ‘allowing’ a site with ω > 1. We next employed branch-site models (test 2) to explicitly test for the rates of evolution on a site along a specific lineage of a tree: branch-site model (Ma) vs. branch-site model with fixed ω1 = 1 (Ma0) (Zhang et al. 2005). The Ma model assumes that sites on the foreground branches are under positive selection. When the LRT was significant (*p* < 0.05) under the M8 and Ma, the codon sites under positive selection were assessed using the Bayes Empirical Bayes (BEB) method (Yang et al. 2005). We only accepted positively selected sites with a posterior probability (PP) > 0.80. A series models implemented in HyPhy (http://www.datamonkey.org) (Pond and Muse 2005) were also used to estimate ratios of nonsynonymous (*d_N_*) and synonymous (*d_S_*) based on a maximum likelihood (ML) framework (Pond and Frost 2005a), including the Single Likelihood Ancestor Counting (SLAC), the Fixed Effect Likelihood (FEL), the Random Effect Likelihood (REL) (Pond and Frost 2005b). As the criteria to identify candidates under selection, we used a Bayes Factor >50 for the REL and a cut-off *P*-value of 0.1 for the SLAC and FEL. The degree of radical changes and positive selection of amino acid were identified by TreeSAAP v3.2, which detects selection based on 31 physicochemical amino acids properties. Functional information was searched from UniProt (https://www.uniprot.org) (Consortium 2015).

### Comparison of mammalian niches

To investigate the potential link between molecular evolution of GST genes and ecological adaptations, we assigned habitats (aquatic vs. terrestrial) according to data in the literature (Uhen 2007). We used Clade Model C (CmC) to identify the level of divergent selection among clade with different ecological niches and to test what partition of clades best fit the data (Bielawski and Yang 2004). CmC allows site variation among priori defined foreground (cetacean, pinnipeds, sirenians) and background partitions (figure S1), which was compared with the M2a_rel null mode that does not allow variation among divisions. Taking into account the phylogenetic relationships between mammals, a phylogenetic ANOVA using the function *phylANOVA* implemented in the R package ‘phytools’ (Revell 2012) was performed to test for differences in the average gene number of functional GSTs between marine and terrestrial mammals. A comparison of GST activities of blood obtained from various terrestrial and aquatic species was obtained from (Wilhelm et al. 2002).

## Results

### Cytosolic glutathione transferase gene repertoires in mammals

To examine the evolution of cytosolic glutathione transferase (GST) genes in mammals we interrogated public genomic data of 21 species, representing all major mammalian taxa. We identified a total of 448 GST genes (333 intact genes, 22 partial genes, and 93 pseudogenes) (table 1). Amino acid sequence identity between GST paralogous was more than 50%, whereas the identity between all subclasses was less than 30% (table S3).

**Table 1.** Overview of cytosolic glutathione transferase (GST) genes in 21 mammals.

| Subclass | Alpha<br>(GSTA) | Mu<br>(GSTM) | Theta<br>(GSTT) | Pi<br>(GSTP) | Zeta<br>(GSTZ) | Omega<br>(GSTO) | Sigma<br>(HPGDS) | Total number<br>of GSTs | Pseudogeniz<br>ation rate |
| --- | --- | --- | --- | --- | --- | --- | --- | --- | --- |
| **Human** | 12(7:5:0) | 5(0:5:0) | 3(0:3:0) | 1(0:1:0) | 1(0:1:0) | 2(0:2:0) | 1(0:1:0) | 25(7:18:0) | 0.28 |
| **Mouse** | 6(1:5:0) | 10(4:6:0) | 3(0:3:0) | 7(0:7:0) | 1(0:1:0) | 2(0:2:0) | 1(0:1:0) | 30(5:25:0) | 0.17 |
| **Naked mole rat** | 4(1:3:0) | 10(0:10:0) | 2(0:2:0) | 2(0:2:0) | 1(0:1:0) | 2(1:1:0) | 1(0:1:0) | 22(2:20:0) | 0.09 |
| **Bottlenose dolphin** | 3(1:0:2) | 5(3:1:1) | 2(1:1:0) | 3(1:0:2) | 1(0:0:1) | 2(0:1:1) | 1(0:1:0) | 17(6:4:7) | 0.35 |
| **Killer whale** | 3(1:2:0) | 5(4:1:0) | 2(0:2:0) | 3(1:2:0) | 1(0:1:0) | 2(0:2:0) | 1(0:1:0) | 17(6:11:0) | 0.35 |
| **Yangtze finless porpoise** | 3(1:2:0) | 5(4:1:0) | 2(1:1:0) | 3(1:0:2) | 1(0:0:1) | 2(0:2:0) | 1(0:1:0) | 17(7:7:3) | 0.41 |
| **Yangtze river dolphin** | 3(1:2:0) | 5(4:1:0) | 2(1:1:0) | 3(1:2:0) | 1(0:1:0) | 2(0:2:0) | 1(0:1:0) | 17(7:10:0) | 0.41 |
| **Sperm whale** | 3(1:2:0) | 5(4:1:0) | 2(0:1:1) | 3(1:2:0) | 1(0:1:0) | 2(0:2:0) | 1(0:1:0) | 17(6:10:1) | 0.35 |
| **Minke whale** | 3(1:2:0) | 5(3:1:1) | 2(1:1:0) | 3(1:2:0) | 1(0:1:0) | 2(0:1:1) | 1(0:1:0) | 17(6:9:2) | 0.35 |
| **Bowhead whale** | 2(0:2:0) | 5(3:1:1) | 2(1:1:0) | 3(1:0:2) | 1(0:0:1) | 2(0:2:0) | 1(0:1:0) | 16(5:7:4) | 0.31 |
| **Cow** | 6(2:4:0) | 8(2:6:0) | 2(0:2:0) | 2(0:2:0) | 1(0:1:0) | 3(0:3:0) | 1(0:1:0) | 23(4:19:0) | 0.17 |
| **Tibetan yak** | 6(1:5:0) | 6(0:6:0) | 3(0:3:0) | 2(0:2:0) | 1(0:1:0) | 3(0:3:0) | 1(0:1:0) | 22(1:21:0) | 0.05 |
| **Sheep** | 7(1:6:0) | 8(2:6:0) | 2(0:2:0) | 2(0:2:0) | 1(0:1:0) | 4(0:4:0) | 1(0:1:0) | 25(3:22:0) | 0.12 |
| **Tibetan antelope** | 6(1:5:0) | 6(0:6:0) | 2(0:2:0) | 2(0:2:0) | 1(0:1:0) | 3(0:3:0) | 1(0:1:0) | 21(1:20:0) | 0.05 |
| **Weddell seal** | 4(1:2:1) | 5(2:3:0) | 2(0:2:0) | 7(1:4:2) | 1(0:1:0) | 2(0:2:0) | 1(0:1:0) | 22(4:15:3) | 0.18 |
| **Pacific walrus** | 5(1:4:0) | 8(2:6:0) | 2(0:2:0) | 6(1:5:0) | 1(0:1:0) | 2(0:2:0) | 1(0:1:0) | 25(4:21:0) | 0.16 |
| **Dog** | 6(1:5:0) | 5(3:2:0) | 2(0:2:0) | 12(6:6:0) | 1(0:1:0) | 2(0:2:0) | 1(0:1:0) | 29(10:19:0) | 0.34 |
| **Horse** | 10(1:9:0) | 6(2:4:0) | 2(0:2:0) | 2(0:2:0) | 1(0:1:0) | 2(0:2:0) | 1(0:1:0) | 24(3:21:0) | 0.13 |
| **Microbat** | 6(0:6:0) | 5(1:3:1) | 4(0:4:0) | 6(5:1:0) | 1(0:1:0) | 2(0:2:0) | 1(0:1:0) | 25(6:18:1) | 0.24 |
| **Florida manatee** | 4(0:4:0) | 8(0:8:0) | 2(0:2:0) | 1(0:1:0) | 1(0:1:0) | 2(0:2:0) | 1(0:1:0) | 19(0:19:0) | 0.00 |
| **Elephant** | 3(0:3:0) | 8(0:8:0) | 2(0:1:1) | 1(0:1:0) | 1(0:1:0) | 2(0:2:0) | 1(0:1:0) | 18(0:17:1) | 0.00 |

The number outside brackets is the number of genes in a subclass, while the numbers in the brackets, separated by a colon, indicate the number of pseudogenes, intact gene, and partial genes. Three distinct orders of aquatic mammals are shadowed.

It has been reported that the low genome coverage can effect gene family analyses by introducing frame shifting errors in coding sequences (Young et al. 2010). In our study we employed genomes with good genome coverage in an effort to minimize this potential bias. Both genome coverage and the N50 length (length at which 50% of total assembly length is covered) of contigs and scaffolds are considered important attributes for assessing a genome assembly. Considering the 13 species sequenced by Illumina sequencing alone in our 20-genome data set, scatter plots of sequence coverage (×) or scaffold N50 versus number of identified GST genes did not reveal a significant positive correlation (figure S2). This suggests that we have identified *bona fide* mammalian GST genes.

### The phylogenetic relationships of cytosolic GST genes

We next wished to classify the 448 cytosolic GST genes into their respective subclass. The phylogenic relationships were reconstructed using maximum likelihood and Bayesian methods. Both topologies yielded similar branch patterns, indicating a reliable tree structure (figure S3). Each GST subclass was clustered into a monophyletic group with high node bootstrap values (e.g., 94 –100% of bootstrapping) (figure S3). However, each subclass across species were not clearly resolved (<50% node bootstrap support) (figure S3); especially for alpha and mu, where duplication were common. According to the phylogenic tree, 105 (81 genes and 24 pseudogenes), 133 (90 genes and 43 pseudogenes), 47 (42 genes and 5 pseudogenes), 74 (54 genes and 20 pseudogenes), 21 (21 genes and 0 pseudogenes), 47 (46 genes and 1 pseudogenes), and 21 (21 genes and 0 pseudogenes) genes across 21 species respectively clustered into alpha, mu, theta, pi, zeta, omega, and sigma subclass, indicating that the cytosolic GST gene family is highly conserved in mammals (figure 1).

**Figure 1.**
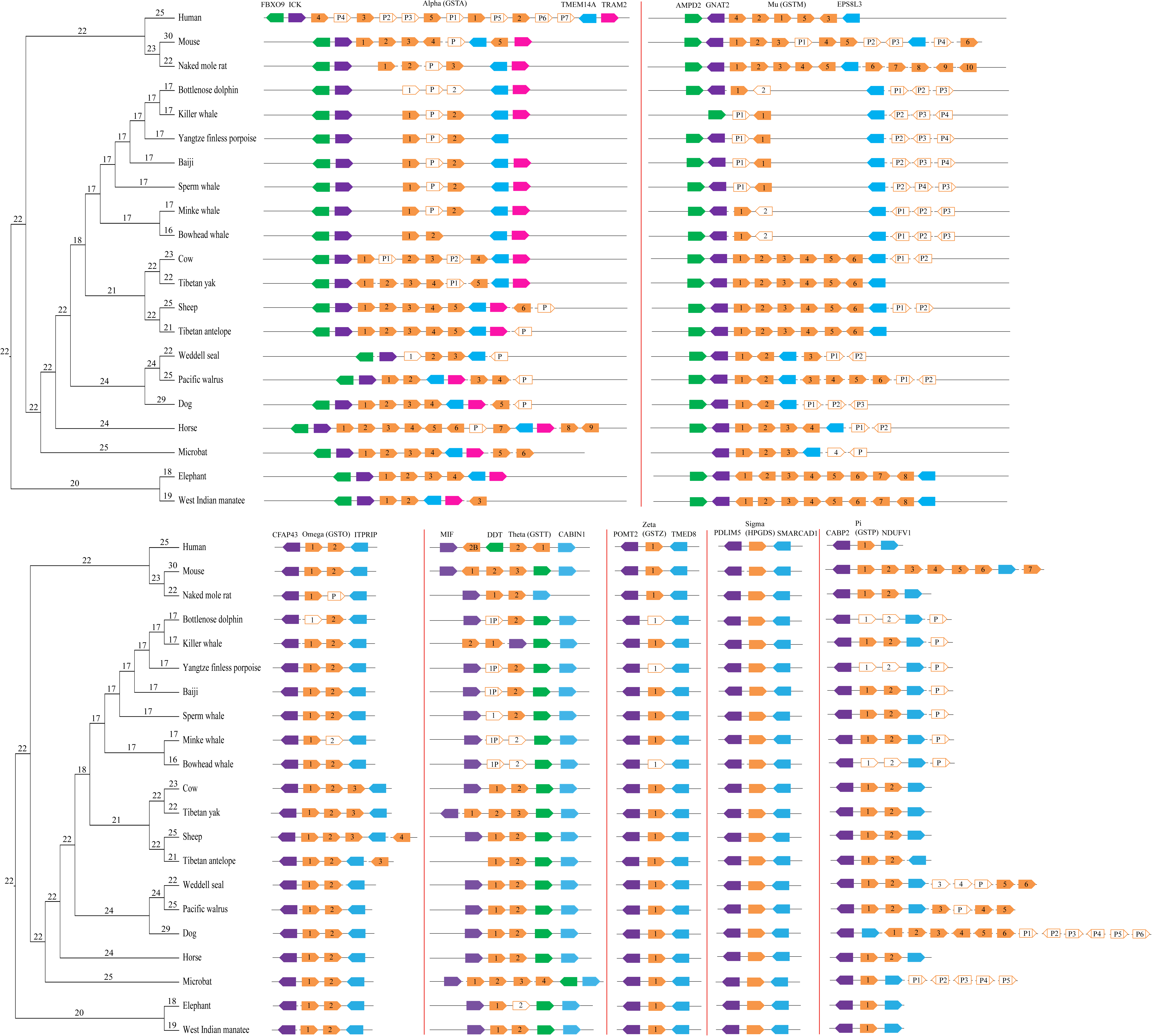
Genomic organization of GST genes in 21 mammalian species according to the phylogenetic tree, genomic position and blast results. Connecting lines indicate GST genes on the same chromosome/genomic scaffold. All GST genes are shown in orange, and the flanking genes are indicated in green, purple, blue and pink. The gene family sizes for ancestral states are shown along each node in the phylogenetic tree. Filled figures: intact genes; empty figures: partial gene, and empty figures with a vertical line: pseudogenes (P).

This scenario was further supported by synteny analysis. All seven cytosolic GST subclasses showed conserved synteny and similar arrangement across mammalian genomes (figure 1). Duplicated genes within subclass were arranged in a tandem cluster, such as members of alpha, mu, theta, pi and omega class, with two or more copies in tandem arrangements. However, the zeta and sigma classes had a single copy per species. Alpha, mu, theta, pi, zeta, omega, and sigma class genes were flanked by the following genes: *ICK* and *TMEM14A*; *GNAT2* and *EPS8L3*; *MIF* and *CABIN1*; *CABP2* and *NDUFV1*; *POMT2* and *TMED8*; *CFAP43* and *ITPRIP*; *PDLIM5* and *SMARCAD1* (figure 1). This further strengthens our GST gene predications. A notable exception was genes of the pi subclass, where genes in the dog where present on separate scaffolds, however we surmise that this could reflect from a sequencing or genome assembly artifact.

Notably, the ML and Bayesian phylogenies based on 448 GST genes did not arrange the seven subclasses into well-supported clades (< 50% node bootstrap support). This could result from the 93 pseudogenes and 22 partial GST genes that are likely no longer under natural selection pressure. Therefore, only the 333 intact (complete CDS) sequences were further used for inferring the phylogenetic tree and determine the evolutionary relationship of subclasses. According this phylogeny, Mu clustered with the pi subclass, but the bootstrap support value and posterior probability were low (BS = 18%; PP = 0.34, figure 2). Alpha grouped with sigma with high bootstrapping and posterior probability (BS = 91%; PP = 0.58, figure 2). The mu, pi, alpha, and sigma subclasses were much more closely related to each other, with 98% bootstrap support and 0.50 posterior probability (figure 2). The theta subclass was placed as sister to a clade containing mu, pi, alpha, and sigma genes with high support (BS = 85%; PP = 1.00, figure 2). Moreover, these five subclasses formed a monophyletic sister clade to the zeta subclass, with 85% bootstrap support and 1.00 posterior probability (figure 2). In addition, the phylogenetic tree recovered the omega subclass as the first-diverging lineage within GSTs (figure 2).

**Figure 2.**
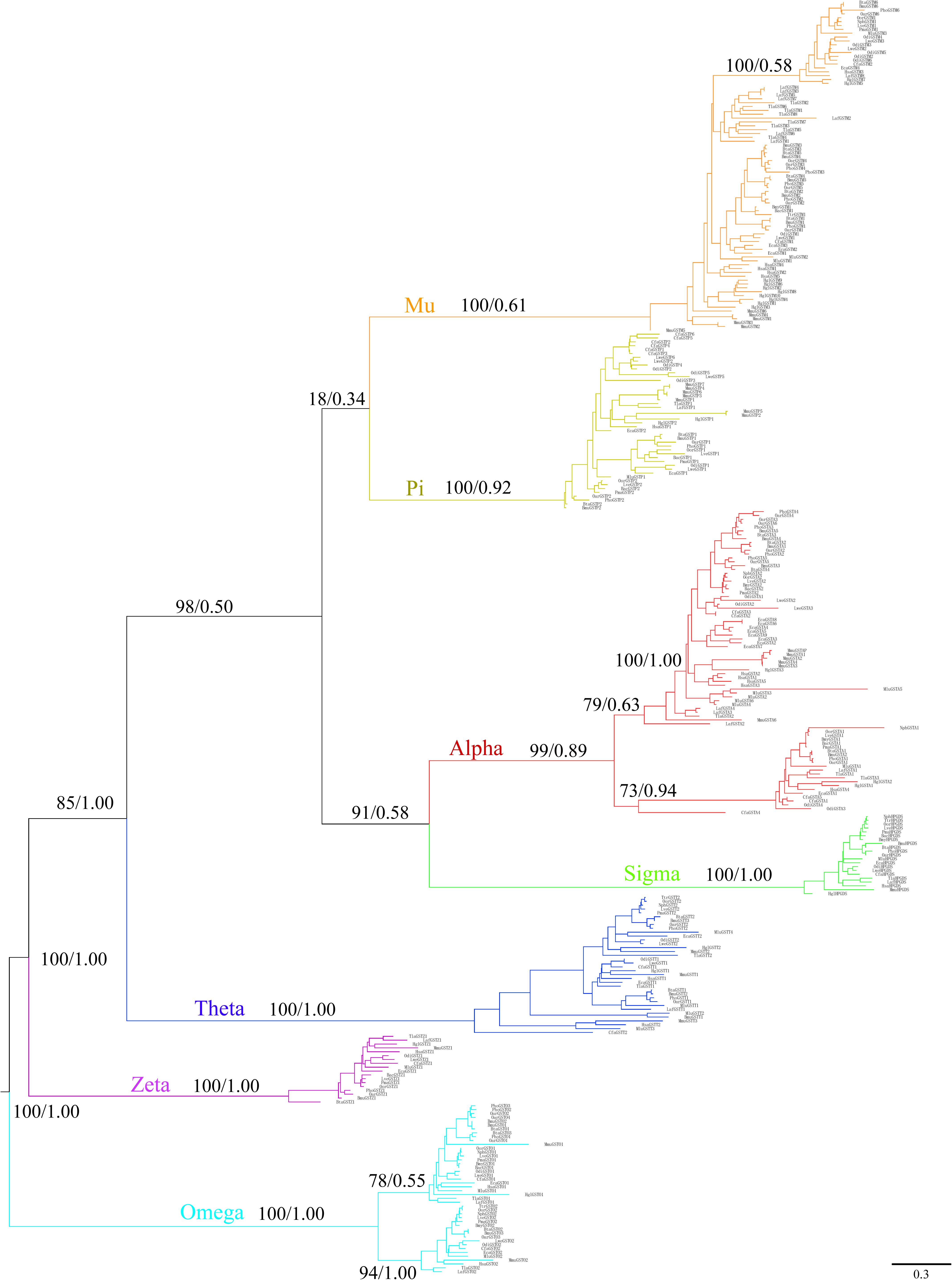
Phylogenetic tree of GST gene family in mammals. Maximum likelihood and Bayesian phylograms describing phylogenetic relationships among the 333 intact sequences of mammalian GSTs. Numbers of the nodes correspond to maximum likelihood bootstrap support values and Bayesian posterior probabilities. The clades of alpha, mu, pi, omega, sigma, zeta, theta are shown in red, orange, turquoise, azure, green, purple, and blue, respectively.

### Lineage-specific gene duplication and deletions

Phylogenetic trees of the GST sequences we identified contain different copy number along mammalian lineages. Of the 21 species studied, mouse (25 intact GSTs), naked mole rat (20 intact GSTs), Tibetan yak (21 intact GSTs), Tibetan antelope (20 intact GSTs), sheep (22 intact GSTs), Pacific walrus (21 intact GSTs), and horse (21 intact GSTs) have the largest GST repertoires (table 1, figure 1). On the opposite spectrum, cetaceans appear to have the smallest GST repertoire – about ten functional GSTs per species (table 1, figure 1). In agreement, the fraction of GST pseudogenes is the highest in cetaceans (mean, 36%, table 1), which is three times higher than terrestrial artiodactyls (mean, 11%, table 1). Considering the alpha subclass, we found that the cetacean alpha subclass consists of two functional GSTs, but their relatives (i.e., artiodactyls) have more than four intact alpha GSTs. Moreover, sheep (6 intact GSTAs), horse (9 intact GSTAs), and microbat (6 intact GSTAs) have very large of gene number in alpha GSTs. Similarly, only two functional mu GSTs were identified in cetaceans. In contrast, the number of functional GSTM genes in artiodactyls (6) is almost six times that of cetaceans. Notably, a large group of species, including all rodents and the two afrotherians, also harbors six to ten functional mu GSTs (table 1, figure 1). Additionally, microbat (4 intact GSTTs) has the largest gene number of theta GSTs, while the lowest gene copy number was found in cetaceans (just one gene in six cetaceans; two in killer whale). Almost all mammalian species have two omega subclass genes, with the exception of artiodactyls (3 or 4 genes) (table 1, figure 1). The zeta and sigma classes appear to be more conserved in mammals, with one copy identified, respectively. Furthermore, the gene gain and loss of GSTs at each ancestral node was estimated by the software CAFÉ. The estimated number of GST genes in the last common ancestor (LCA) of cetaceans (17) is considerably smaller than that in the LCA of artiodactyls (21) (figure 1). Moreover, the LCA of carnivorans and rodents was estimated to have a larger repertoire of GST genes than in the remaining lineages.

### Evolutionary model of cytosolic GST gene family in mammals

To investigate the possible role of natural selection in the evolutionary history of cytosolic GST gene family, a series of site-specific and branch-specific models were conducted. Site-specific selection tests implemented in PAML were performed to assess the selective pressure acting on mammalian GSTs. GSTA1, GSTM1, GSTO1, GSTO2, GSTP1, GSTP2, GSTT2, and GSTZ1 showed site-specific positive selection; with 16, 5, 14, 3, 2, 6, 1, and 3 sites having posterior probabilities > 0.80 (table S4); suggesting that GSTs evolved under diversifying selection in mammals. Similarly, 23, 28, and 27 positively selected codons in eight genes were identified by SLAC, FEL, and REL model implemented in Datamonkey, respectively. In total, 38 codons from eight genes (*GSTA1*: 10; *GSTM1*: 1; *GSTO1*: 9; *GSTO2*: 5; *GSTP1*: 2; *GSTP2*: 5; *GSTT2*: 3; and *GSTZ1*: 3) with evidence of positive selection were detected with at least two of the ML methods (table 2). Among these codons, 31 sites were also detected by amino-acid level selection analyses using TreeSAAP. Six of these sites possess more than six radical amino acid changes (table 2).

**Table 2.**
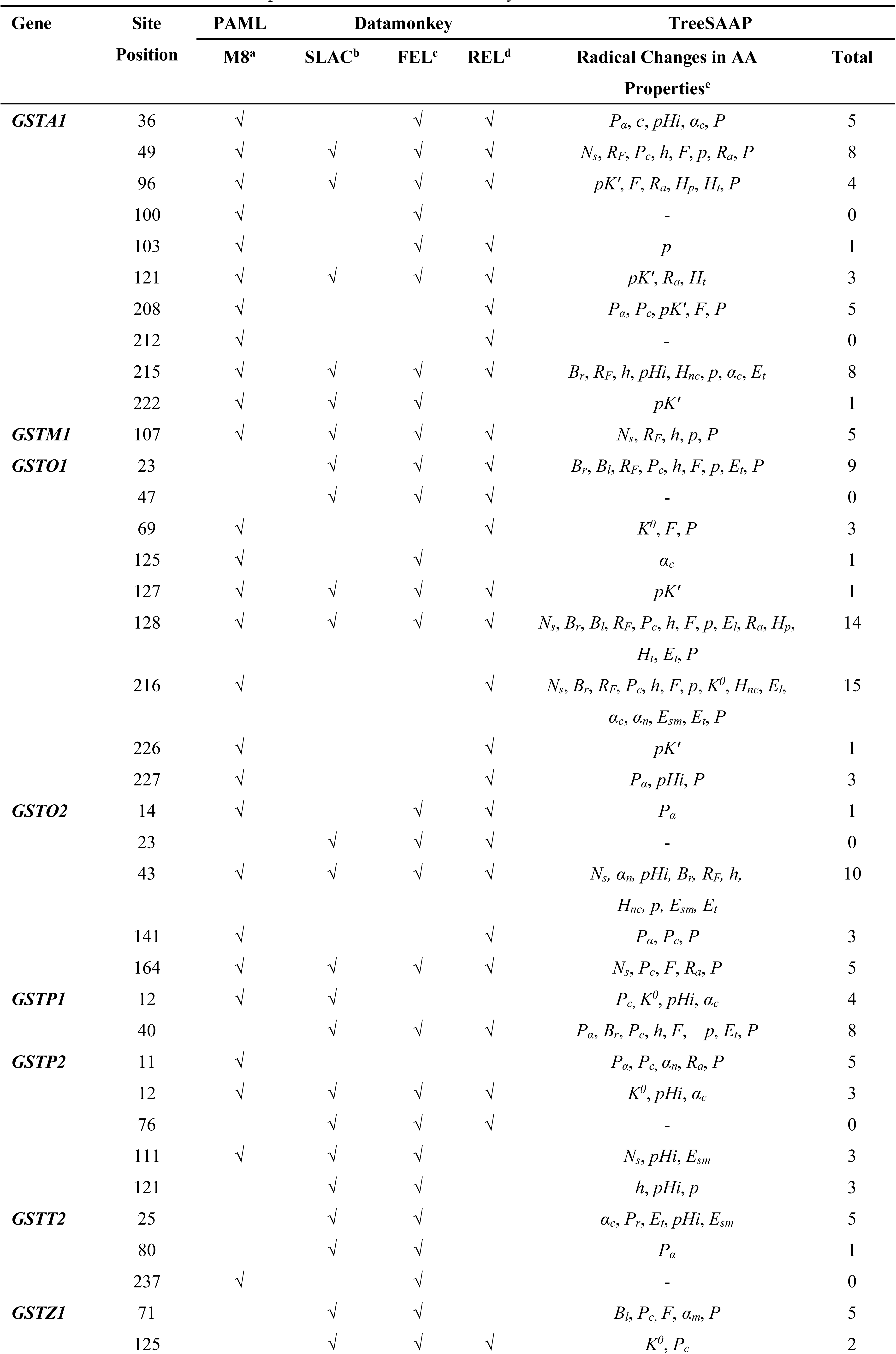
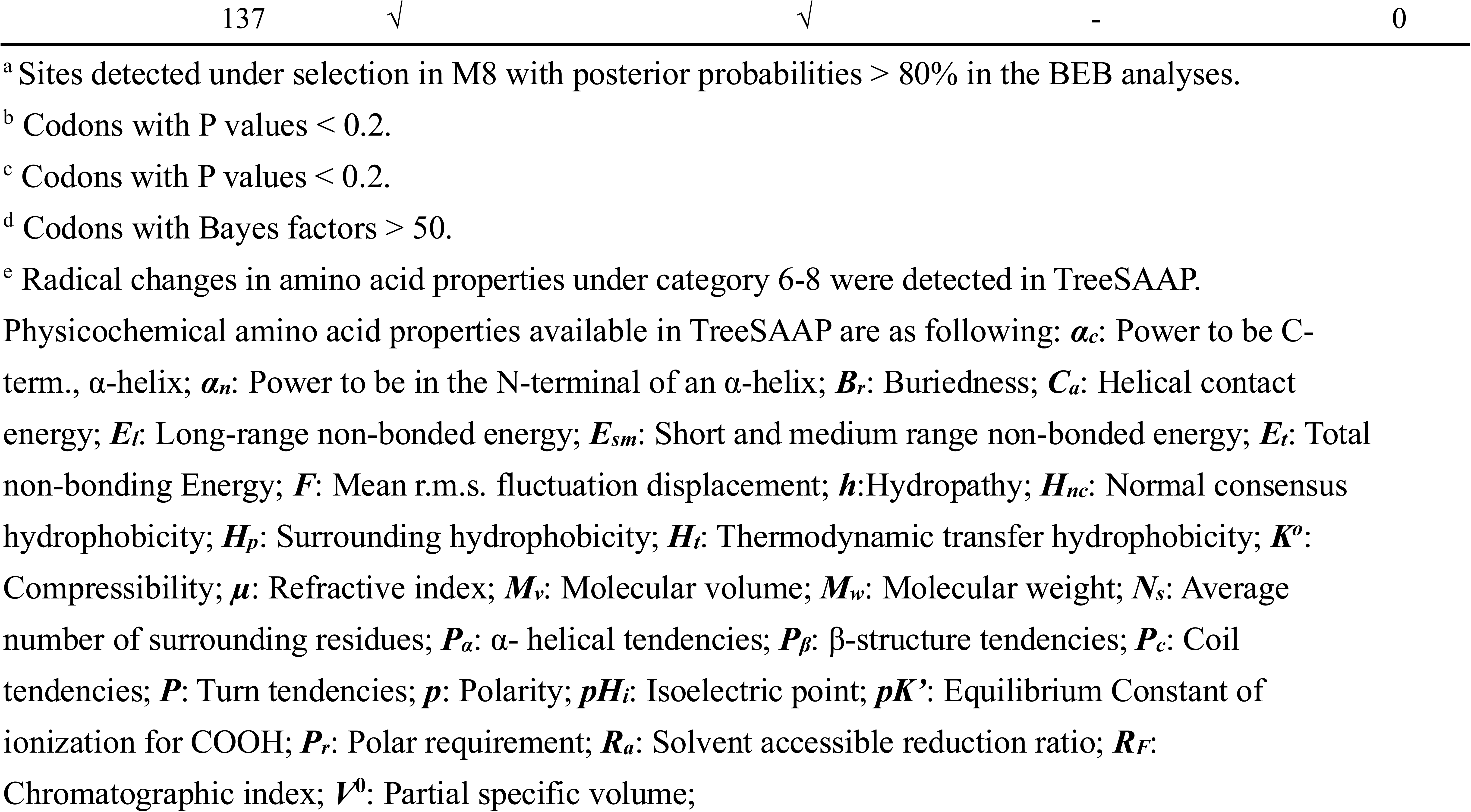
Amino acid sites under positive selection detected by ML methods

We further employed the branch-site model to identify the episodic adaptations that affect amino acids along specific lineages. A few of internal branches and several terminal branches showed evidence of positive selection after correction for multiple testing (table 3, figure S4). In cetaceans the lineage leading to *GSTO1* in bottlenose dolphin and *GSTP2* in sperm whale were under selection. In pinnipeds the branches leading to *GSTM1* in Pacific walrus and the ancestor, *GSTP2* in the ancestor, as well as *GSTA1* in Weddell seal were under strong positive selection. *GSTA1* and *GSTM1* detected to be under positive selection only in pinnipeds (here: Pacific walrus and Weddell seal). Likewise, the lineage leading to human *GSTA4*, elephant, and microbat *GSTT1*, cow, horse and manatee *GSTT2*, sheep and antelope *GSTP2*, naked mole rat *GSTO1*, as well as cetartiodactyla (includes whales and dolphins, and even-toed ungulates) HPGDS were also under positive selection.

**Table 3.**
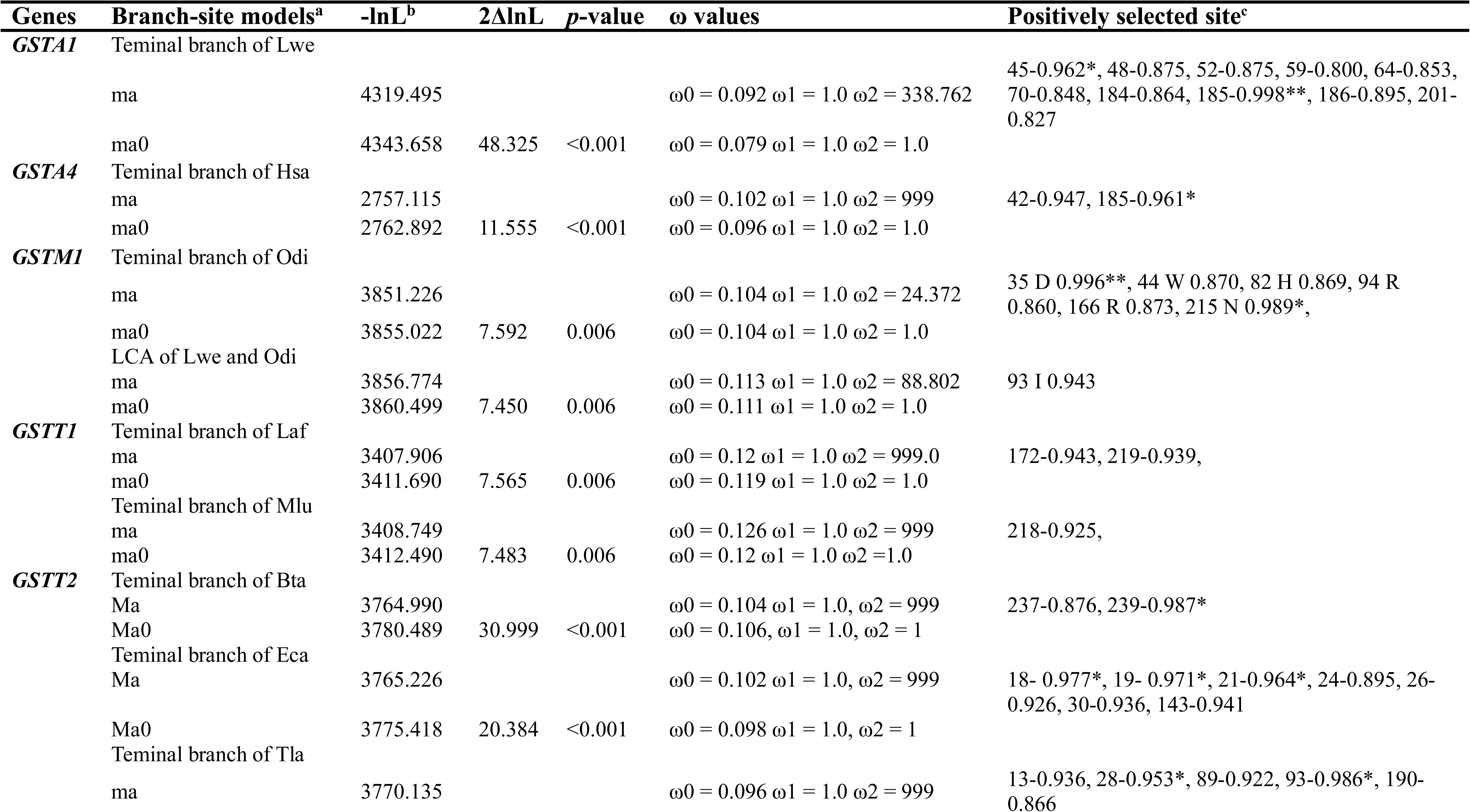
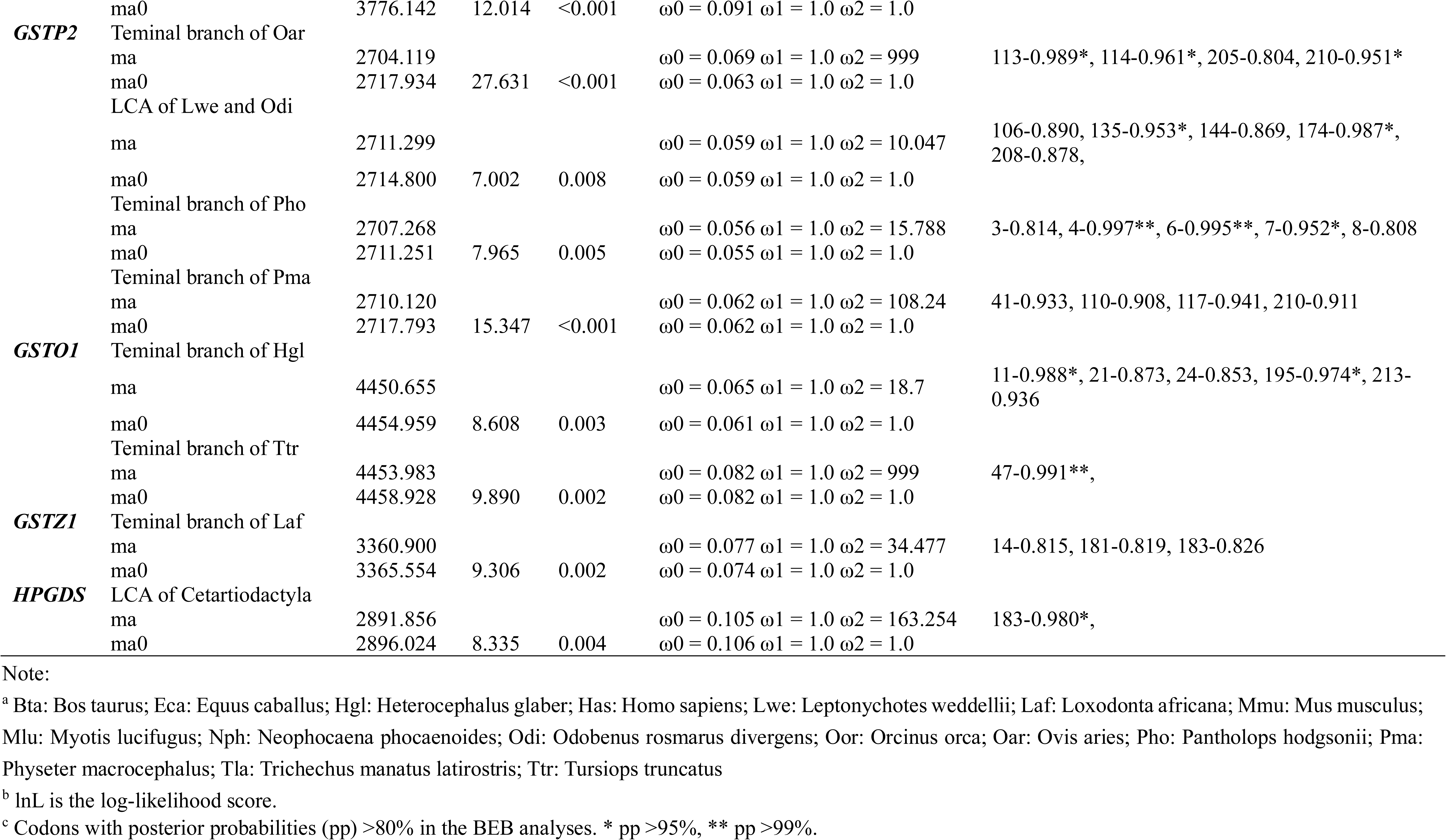
Selective pattern analyzed by Branch-site model

### Varied gene family numbers and enzyme activities of cytosolic GSTs according to habitat

A more thorough investigation of the evolutionary history of the mammalian GSTs was conducted by extending the analysis to divergent ecological groups (marine vs. terrestrial). The clade model C, which assumes divergent selection along the cetacean, pinniped, sirenian, and marine mammal branches fitted the data better than the null model in the case of the GST genes *GSTA1*, *GSTA4*, *GSTM1*, *GSTM3*, and *GSTT1* (*P* < 0.05, table S5), suggesting there is a divergent selection pressure between marine and terrestrial mammals and within marine mammal groups. Moreover, the two-partition that divided cetaceans/pinnipeds and sirenians was a significantly better fit than the one-partition marine mammal group in the case of *GSTA1* (table S5). The best three-partition model was found to be the division of cetacean, pinniped, and sirenian at the *GSTA1* gene (table S5).

To explicitly test whether ecological niches affect the number of functional GST genes and the molecular evolution of GSTs, we performed several statistical tests selecting ecological factor (marine vs. terrestrial) as predictor variables. We found that the gene number of all GSTs (phylANOVA; *F* = 23.135, *p* = 0.009) and alpha class GSTs (phylANOVA; *F* = 21.599, *p* = 0.007) were significantly influenced by ecological factors. The total number of GSTs and alpha subclass gene number along marine mammals were significantly smaller than terrestrial species (figure 3A). Next, we further investigated the evolutionary dynamics of GST genes among divergent groups (i.e., cetacean, pinniped, sirenian) separately. There was no significant difference between individually aquatic group and terrestrial species except for cetaceans. Of note, cetaceans had a smaller cytosolic GST repertoire (phylANOVA; *F* = 142.458, *p* = 0.001) than terrestrial mammals (figure 3B). Moreover, the result showed that the estimated number of alpha class GSTs (phylANOVA; *F* = 24.300, *p* = 0.026) and mu GSTs (phylANOVA; *F* = 27.557, *p* = 0.011) in cetaceans is also lesser than those in terrestrial mammals.

**Figure 3.**
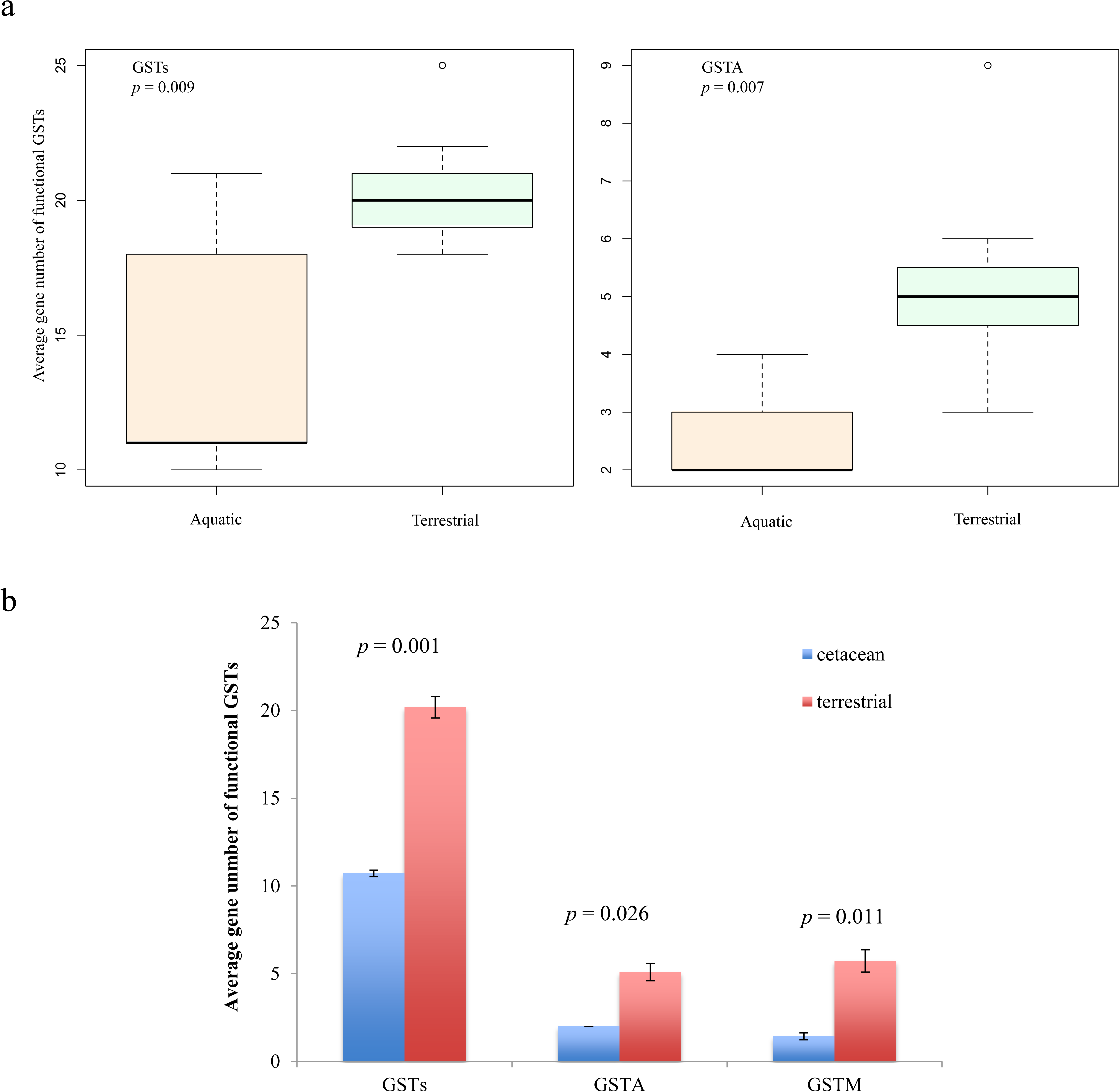
Numbers of functional GST families were compared across ecological niches using phylANOVA. a) aquatic vs. terrestrial mammals; b) cetacean vs. terrestrial species.

A comparison of publicly-available data on blood GST enzyme activity from aquatic and terrestrial mammals was examined (table 4), indicating that GST activity of pinnipeds and sirenians are generally higher than that of terrestrial relatives (species in Carnivora and Xenarthra). However, the GST activity of cetaceans is lower compared to artiodactyls. Among marine mammals, sirenians has the highest enzyme activities of, followed by pinnipeds; while the lowest GST activity was found in cetaceans (figure 4).

**Table 4.** Enzyme activity of glutathione S-transferase (GST) in blood of marine and terrestrial mammals.

| Order | Common name | Latin name | GST ($\mu\text{mol ml}^{-1} \text{min}^{-1}$) |
| --- | --- | --- | --- |
| Marine mammals |  |  |  |
| Cetacea | Minke whale | *Balaenoptera acutorostrata* | 276.04 |
| Cetacea | Striped dolphin | *Stenella clymene* | 139.06 |
| Sirenia | Florida manatee | *Trichechus manatus manatus* | 588.97 |
| Pinnipedia | Southern elephant seal | *Mirounga leonina* | 267.29 |
| Terrestrial mammals |  |  |  |
| Artiodactyla | Pampas deer | *Ozotocerus bezoarticus* | 292.19 |
| Xenarthra | Southern tamandua | *Tamandua tetradactyla* | 228.05 |
| Carnivora | Grison | *Grison vittatus* | 280.49 |
| Carnivora | South American racoon | *Procyon cancrivorous* | 142.95 |

**Figure 4.**
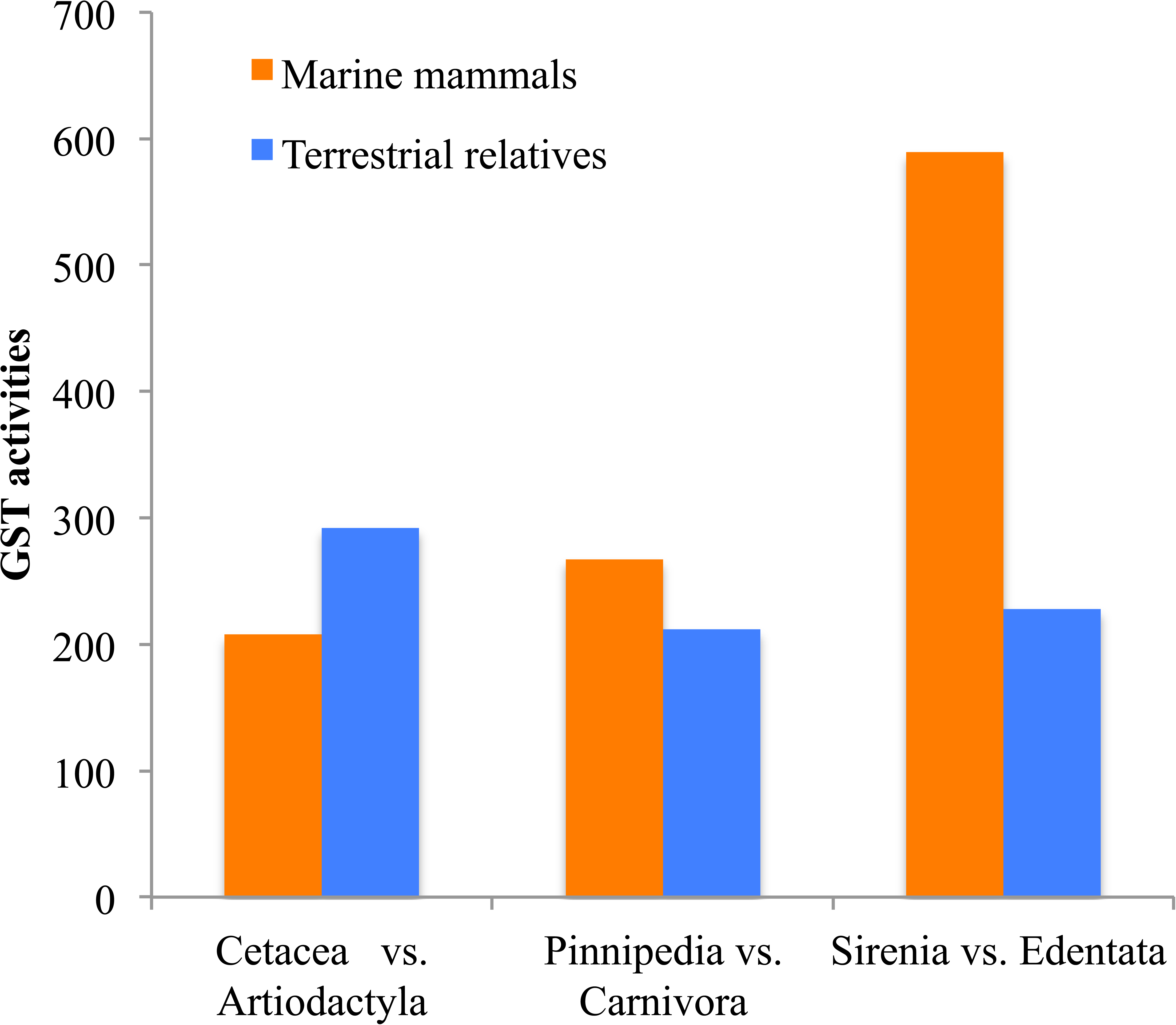
Histogram showing the comparison of GST activities in blood between marine mammals and their relatives.

## Discussion

### Molecular evolution of GST genes

Several previous surveys have documented the distribution of GST. Nebert and Vasiliou provided a comprehensive assessment of GST superfamily genes (Nebert and Vasiliou 2004). Other studies examined the phylogeny (Pearson 2005) and evolution (da Fonseca et al. 2010) of the superfamily across the tree of life. In the present study, we expanded on the previous surveys by performing a comprehensive search for cytosolic GST gene 20 mammals representative of all major mammalian taxa. Of these, 15 species had no previous information on their cytosolic GST gene repertoire (Figure 1). We provide evidence for positive selection acting on several GST genes in divergent taxonomic groups (*GSTA1*, *GSTM1*, *GSTO1*, *GSTO2*, *GSTP1*, *GSTP2*, *GSTT2*, and *GSTZ1*), indicative of that pervasive adaptive evolution. Considering that cytosolic GST gene play critical roles in the detoxification and metabolic activation of xenobiotics (Board and Menon 2013), these changes are likely to reflect previous and ongoing adaptions to diverse environments by mammals. In addition, 31 positively selected sites with radical amino acid changes were identified by gene and protein-level selection analyses, indicating that a single amino acid change can alter the specificity or potency of detoxification and antioxidant defenses during mammalian evolution. In particular, a series of positively selected sites were found to be located in or close to functional domains. For example, site 45 in GSTA1 is known to be involved in GSH binding (Balogh et al. 2009), while GSTA1 sites 212 and 215 are close to the site 216 important for thiolester substrate hydrolysis (Hederos et al. 2004). Additionally, His107 in GSTM1 is a second substrate-binding site and site-directed mutation experiments (His107Asn) revealed a 50% reduction in catalytic activity (Patskovsky et al. 2006). Thus, it is plausible that the positively selected sites identified contribute to altered GSTs enzyme.

### Mammalian cytosolic GSTs phylogeny and birth-and-death evolution

The phylogenetic relationships among 333 cytosolic GSTs were reconstructed by two independent methods which gave a similar estimation of mammalian phylogeny. The branch pattern indicated that the omega, theta, and zeta subclass are ancient in mammals, while the alpha, mu, pi and sigma subclasses evolved later (figure 2). This is in agreement with previous studies based on structural and functional data (Armstrong 1997; da Fonseca et al. 2010; Frova 2006). The omega, theta, and zeta subclass employ catalytic serine hydroxyl to activate GSH, and they are thus predicted to be the progenitors of GSTs (Frova 2006). According to our phylogenetic tree, the omega subclass harbors the most ancient genes (figure 2). Omega GSTs have strong homology to glutaredoxins, the predicted ancestors of N-terminal topology of GST (Frova 2006; Oakley 2005). Taken together, it is thus reasonable to assume that the omega subclass evolved earlier. A switching from serine to tyrosine in the alpha, mu, pi, and sigma subclasses is another evolutionary scenario of GSTs, which would explain why these four subclasses cluster together in the tree with high bootstrapping (figure 2). It would appear that the sigma subclass diverged before the mammalian alpha, mu, and pi group due to it presents in invertebrates and vertebrates (Frova 2006). In the present case, however, sigma appeared as the sister group of the alpha subclass, which is in line with a previous study (da Fonseca et al. 2010). The mammalian sigma subclass, known as prostaglandin synthase, has a hydrophilic interface with a lock-and-key motif – similar to the alpha, mu, and pi subclasses (Sheehan et al. 2001). Therefore, sigma clustered with alpha subclass might be related to their specialized structure and function in mammals. It is interesting to note that heterodimers can be formed between subclass mu and pi polypeptides, reflecting close evolutionary relationships (Pettigrew and Colman 2001). This suggestion was further supported by the clustering between subclass mu and pi in our phylogenetic tree, although the bootstrap support values and posterior probabilities were low. On the other hand, this result also suggests that subclass mu proteins arose most recently. The theta subclass was placed as sister to a clade containing mu, pi, alpha, and sigma subclass with high support, supporting the prediction that alpha, mu, pi, and sigma subclasses likely arose from the duplication of theta subclass (Armstrong 1997).

The GST gene family has been previously described in prokaryotes and eukaryotes (Pearson 2005), reflecting different expansion and contraction. In this study, we examined cytosolic GST gene repertories in 21 species representing all major mammalian taxa. Our results show an unequal copy number of GST genes across the mammalian phylogeny, as it has previously been suggested (da Fonseca et al. 2010; Pearson 2005). For instance, the largest gene expansion was observed in the mouse (30 copies), whereas only 16 GSTs were identified in bowhead whale. This could be due to a lineage or species-specific duplication or deletions of this gene family in mammals. In support of this possibility, a diverse pseudogenization rate was further found in mammalian GSTs; ranging from 41% in Yangtze river dolphin and Yangtze finless porpoise to 0% in elephant and Florida manatee (table 1). Studies of gene duplicates show that new genes are created by gene duplication and some duplicated genes maintain in the genome for a long time, nonetheless others are nonfunctional or deleted from the genome. This phenomenon is termed the birth-and-death evolution model (Lynch and Conery 2000; Nei and Rooney 2005). This model is well supported by the gene duplication events observed in our data of mammalian GSTs.

It is also important to note that extensive duplication (12 paralogous) of pi GSTs was found in the dog, whereas 50% of them were pseudogenization events (human alpha subclass in the same case). It has been reported that most duplicated genes tend to experience a brief period of relaxed selection early in their history, which result in non-functionalization or pseudogenes (Lynch and Conery 2000). Therefore, the high fraction of pseudogene in dog pi GSTs (or human alpha GSTs) further supports the birth-and-death model of GSTs evolution. Notably, however, seven pi subclass paralogous identified in mouse are apparently intact and functional, as is the case for the naked mole rat mu GSTs (10 copies), which could be the outcome of adaptations to environmental substrates via diversification of duplicated copies. Moreover, our results revealed extensive positive selection in *GSTP2* along five mammalian lineages (figure S4), suggesting of episodic selection pressure – probably in response to changes in xenobiotic exposure. In contrast, there is no evidence of positive selection in *GSTP1* along specific lineages, indicating a functional conservation which is consistent with the critical role of GSTP1 in ethacrynic acid metabolism in the liver (Henderson et al. 1998). These results reveal that divergent selective regimes occurred in paralogs within a cytosolic GST class.

### Divergent selection in mammalian cytosolic GSTs and ecological niche

Some of our striking results are the presence or absence of one or several cytosolic GST genes between species living in distinct ecological milieus (table 1). For example, in carnivores, 29 genes were identified in dog, whereas 22 and 25 genes were found in the Weddell seal and Pacific walrus, respectively. In cetartiodactyla, 21 to 25 genes can be found in artiodactyla, while cetaceans possess only 16 or 17 genes. Numerous studies have found that gene family evolution is closely tied with environments. This includes opsin genes in cichlid fish (Henderson et al. 1998), hemoglobins in vertebrates (Nery et al. 2013), keratin-associated proteins (Khan 2014; Sun et al. 2017), and olfactory receptor genes in mammals (Hughes et al. 2018; Niimura et al. 2018). We, therefore, hypothesized that dynamic evolution of GST gene family is related to ecological adaptations in aquatic and terrestrial species. As expected, we demonstrated that the number of GST genes is significantly correlated with the habitat, suggesting that hypoxia-tolerant species tend to have fewer GST genes (figure 3a). Coincident with contrasting gene number, moreover, our analysis revealed that hypoxia-tolerant species with different ecological niches showed evidences of divergent selection (table S5), further suggesting that adaptive evolution related to habitats could play a role.

Interestingly, we found that the number of cytosolic GST genes in cetaceans was significantly less than terrestrials (figure 3b), suggesting gene loss along the cetacean branches played a major role. It has been reported that genes expressed at lower levels tend to be more prone to loss in lineages (Krylov et al. 2003). Considering the plausibility that expression levels determine the enzyme activities of the tissues (Tiedge et al. 1997; Rodríguez-Antona et al. 2001), our results of reduced GST genes and lower GST enzyme activity in cetaceans possibly support this prediction (figure 3b, 4).

The contraction of alpha GSTs is striking, with only two functional GSTAs (*GSTA1* and *GSTA4*) in cetaceans. Similarly, five mu GSTs were identified in cetaceans, but only one intact gene (GSTM1) keeps its function compared with 5-10 functional GSTMs in other mammals (figure 1). Contraction of the cetacean GSTs appears to have occurred after the divergence of cetacea from artiodactyla approximately 53 Mya (million years ago) (Thewissen et al. 2007), undoubtedly reflecting many of the unique adaptations of the aquatic lifestyle. The presence of four GSTM pseudogenes provides further evidence for the contraction of the cetacean locus from a large GST family in the ancestral cetacean genome. Cetaceans are continually subjected to oxidant challenges not only because of chemical pollutants in aquatic environment, but also due to the hypoxia/reoxygenation or ischemia/reperfusion processes which induces free radical reactions, production of toxic reactive oxygen species (ROS), and increased superoxides (e.g., xanthine and xanthine oxidase) thus leading to oxidative damage in a wide variety of tissues (Valavanidis et al. 2006; Li and Jackson 2002). The pseudogenes in cetaceans showed a signature of relaxed functional constraint (table s6), suggesting that cetaceans have degenerated antioxidant defenses; however, they were not compromised because the most critical GSTs are nevertheless retained in cetaceans, showing a constitutive strategy to prevent oxidative damage. For example, *GSTA1* possess a wide range of catalytic properties, which would be expected to be the prime site of metabolism of xenobiotics (Coles and Kadlubar 2005). Again, it has been implicated that GSTA4 plays a vital role in against lipid peroxidation products, induced by the accumulation of ROS during oxygen deprivation stress (Hayes et al. 2005). GSTM1 appears to increase the specific activity approximately five fold toward CDNB via stimulation of superoxide and ROS (Hayes and Pulford 1995). On the other hand, the peroxiredoxin (PRDX) and glutathione peroxidase (GPX) family that eliminate peroxides and protect against oxidative damage in the antioxidant system are expanded in whale lineages (Yim et al. 2014; Zhou et al. 2018), a phenomenon which may indicate compensation for the loss of GSTs. Additionally, we also observed the evidence for positive selection on GSTs in cetaceans, confirming to our speculation that widely dispersed xenobiotics in aquatic ecosystems and high oxidative stress might drive the adaptive evolution of retained functional GST genes in cetaceans to avoid the damages. Our results are consistent with the ‘less-is-more’ hypothesis (Olson 1999). That is, cetaceans employ fewer GST genes to efficiently perform the functions of more GSTs in other mammals and have a strategy that is constitutively ready to respond to oxidative stress.

The aquatic mammals (cetaceans, pinnipeds and sirenians) encompass phenotypic convergences that accommodate the challenges of aquatic living. They present a similar respiratory and cardiovascular solution to tolerant hypoxia and prolong deep dives, such as increased oxygen storing (Ramirez et al. 2007). This is an intriguing scenario the marine mammals facing similar oxidative pressures did not present a similar evolutionary pattern of GST gene family. For instance, heterogeneity of GST gene number and GST activities were found between marine mammals. Beside, 4, 3, and 1 positively selection genes respectively identified along cetaceans, pinnipeds and sirenia (figure S4) also indicates a difference in the strength of positive selection between marine mammals. There are several non-mutually exclusive hypotheses to explain this pattern: 1) Antioxidant status is probably directly related to the diving capacity of diving mammals. Marine mammals that perform shallow/short and deep/long divers might experience different oxidative pressure, suggesting distinct mechanisms to successfully maintain redox balance (Cantú-Medellín et al. 2011); 2) Pinnipeds and sirenians possess enhanced enzymatic antioxidant capacity, whilst non-enzymatic antioxidant (e.g., levels of glutathione) seems to play an important role in cetaceans, indicating a different strategy for antioxidant defenses adaptation (Ninfali and Aluigi 1998; Wilhelm et al. 2002).

## Acknowledgement

This work was financially supported by the National Key Program of Research and Development, Ministry of Science and Technology of China (grant no. 2016YFC0503200 to G.Y. and S.X.), the Key Project of the National Natural Science Foundation of China (NSFC) (Grant no. 31630071 to G.Y.), the National Natural Science Foundation of China (NSFC) (grant nos. 31570379, 31772448 to S.X., grant no. 31872219 to W.R.), the Priority Academic Program Development of Jiangsu Higher Education Institutions (PAPD), and the China Postdoctoral Science Foundation (grant no. 2018M642278 to R.T.).

